# Self-agency built with sensorimotor processing: Decoding self-other action attribution in the human brain

**DOI:** 10.1101/483420

**Authors:** Ryu Ohata, Tomohisa Asai, Hiroshi Kadota, Hiroaki Shigemasu, Kenji Ogawa, Hiroshi Imamizu

## Abstract

A sense of agency can be defined as a subjective experience that I am the one who is causing or generating an action. Several brain regions have been proposed as neural substrates of the subjective experience; however, how the information is processed and organized by each region to achieve the sense of agency still remains unclear. In this study, we have clarified the neural representations corresponding to three processes namely, sensorimotor error, feeling of agency, and judgment of agency. Specifically, we found that the widespread sensorimotor areas represent sensorimotor error information. The right inferior parietal lobe represents the information solely on self-/other-attribution even during movements, which corresponds to the feeling of agency. Finally, the right inferior frontal gyrus shows a distinct representation between self- and other-attribution immediately before reporting the judgment on the movement attribution. These results suggest that the brain builds a sense of agency by developing distinct types of information each corresponding to the three processes with the passage of time.

## Introduction

How the brain makes us aware of our self, as an individual separate from other individuals is a long-standing question in the field of neuroscience. According to William James^1^, a famous philosopher and psychologist, two aspects of self-consciousness should be discriminated; the ‘I’ and the ‘Me’, which can be defined as experiencing oneself as a subject and as an object of perception respectively. The concept of the ‘I’ focuses on the interaction between body and environment (i.e., sensorimotor process) as a process to build up the self as a subject of action and perception^2-4^. While the building-up process represents a fundamental aspect of self-consciousness, studies in cognitive neuroscience have paid little attention to the self as a subject, as compared to the self as an object^5-9^.

The sense of agency, which is one of the representatives of the ‘I’, can be defined as a subjective experience that ‘I’ am the one who is causing or generating an action^10,11^. Neuroimaging studies have reported the involvement of different brain regions in the processes underlying the sense of agency^12-17^. A common strategy among the studies was to subtract the neural activities in the self-attribution condition from the other-attribution condition, and vice versa, while experimentally manipulating a relationship between an action and its outcome. These studies revealed that multiple brain regions, such as the front-parietal network regions are involved in action attribution (review^18,19^). However, what information each region processes and how the brain organizes the information of multiple regions to make a unified decision on self-/other-attribution still remains unclear.

Theoretical models provide a clue for understanding the multiple neural processes underlying the sense of agency. For example, a comparator model, one of the influential models of the sense of agency, explains the mechanism behind incorporation of sensorimotor processing into an agency-attribution system (Fig. 1 left). The brain compares predicted and actual sensory consequences of an action^20-22^. The result of the comparison, known as a prediction error, is incorporated into the system to judge action attribution^21,23^. While the comparator model mainly deals with the sensorimotor process, Synofzik et al.^24^ highlighted the existence of the process corresponding to non-conceptual feeling of agency. This process takes over the outcome of the comparator model and provides the information to the next conceptual judgment process for agency attribution (i.e., judgment of agency, Fig. 1 right). Taken together, the theoretical models emphasized that the sense of agency is formed by a hierarchical system that includes multistep processes ranging from sensorimotor to cognitive processing (Fig. 1). In the current study, we hypothesized that different brain regions are involved in representing information separately for sensorimotor, non-conceptual feeling and conceptual judgment processes.

**Figure 1:**
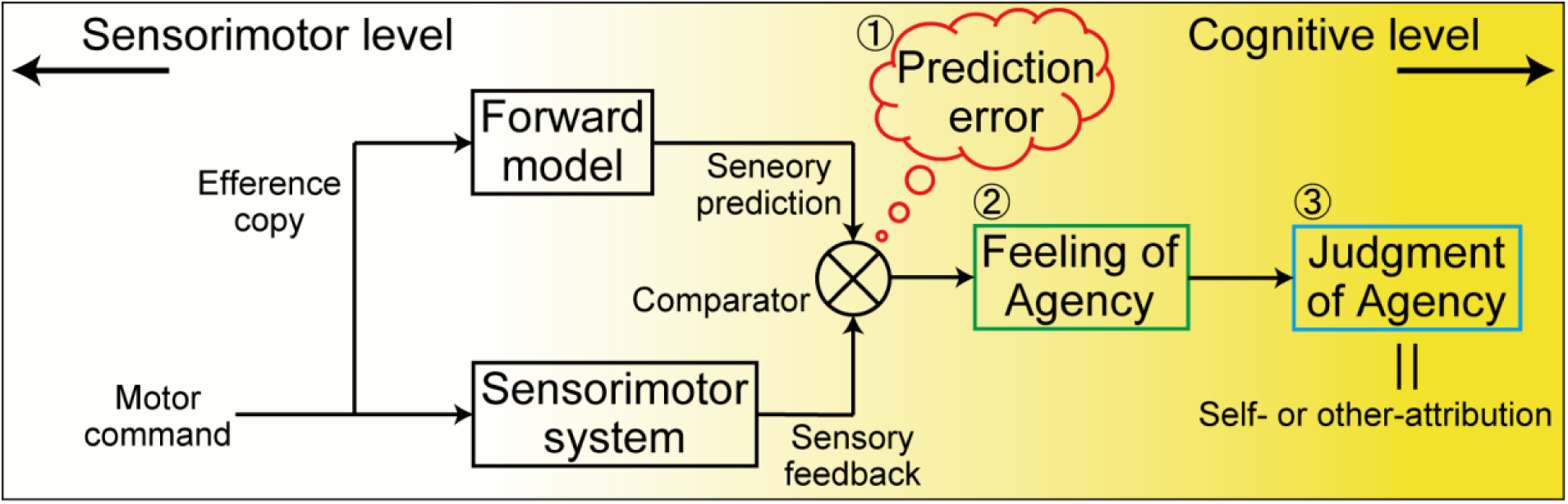
A hierarchical system to form a sense of agency. Theoretical models emphasize that the sense of agency arises through multiple steps from sensorimotor to cognitive level. This schematic represents an overview combing the two most influential theories of the sense of agency; (1) the comparator model^11,21,23^(left) and (2) the two-step account of agency^24^ (right). Accumulation of sensorimotor information, which is mainly prediction error, contributes to experiencing the feeling of agency. Through the above process, we can achieve the judgment of agency, which is equivalent to self- or other-attribution in our experiment. The background color depicts gradation from sensorimotor to cognitive level.

Here, we tested our hypothesis by combining the two strategies. First, we developed a behavioral experimental paradigm in which self-agency gradually evolved over time^25-27^ Participants continuously traced a target path under ambiguous conditions about agency so that they could gradually realize whether sensory feedback was attributed to self- or other-control. This task enabled us to dissociate the individual multistep process within a single paradigm (Fig. 1). The second strategy was multi-voxel pattern analysis (MVPA) of functional magnetic resonance imaging (fMRI) data^28-30^. MVPA makes it possible to explore neural information represented in distinct patterns of fMRI voxel signals^31,32^. In the current study, we decoded separately self-/other-attribution and information about prediction error to clarify how the distinct information was represented in the brain. Thus, combining the above task paradigm and MVPA, we expected to observe specific neural representation corresponding to sensorimotor error, feeling- and judgment-of-agency processes over time. Our results revealed that 1) the widespread area of the sensorimotor and parietal regions represented prediction error information, 2) the right inferior parietal lobe (IPL) contributed to the feeling of agency, and 3) the right inferior frontal gyrus (IFG) and ventrolateral prefrontal cortex (VLPFC) are responsible for the judgment of agency. Collectively, our study demonstrates the specific role of the individual brain regions in the hierarchical system to build up a sense of agency.

## Results

Inside the fMRI scanner, 18 participants traced a five-cycle sinusoidal target with a cursor on a screen by controlling a joystick (Fig. 2A). Data from seven-participants were eliminated from the following analysis to ensure homogeneity of action attribution process (see Supplementary Results for more detailed description on participant’s rejection for fMRI analysis). They were instructed to move the cursor at a half-cycle of a wave in 1 s (i.e., 0.5 Hz) and complete a single continuous movement within the 10-s Move period. In each trial, we morphed visual feedback (i.e., cursor position (*X*, *Y*)) by incorporating the joystick position (*x*’, *y*’) of the other person into the participant’s own joystick position (*x*, *y*) according to the morphing ratio (*α*) (Fig. 2B). Across trials, the ratio, self 0% (other 100%, α = 0) to self 100% (other 0%, α = 1.0) with every 25% step, was randomly varied (Fig. 2C). We instructed the participants to control the cursor to trace the target path as precisely and as smoothly as possible even if the movement of the other person was felt strongly to determine the cursor position. After the tracing, participants were judged within the 8 s rate period about how much they felt the cursor movement was attributed to their own joystick movement on a 9-point Likert scale. The scale ranged from a score of 1 (movement of some other person) to a score of 9 (movement derived completely from the participant). We found a significant effect of morphing ratio on the self-other rating score (*F* (4, 50) = 152.4, *p* < 0.001, see Supplementary Results: “Behavior results” and Supplementary Fig. S1).

**Figure 2:**
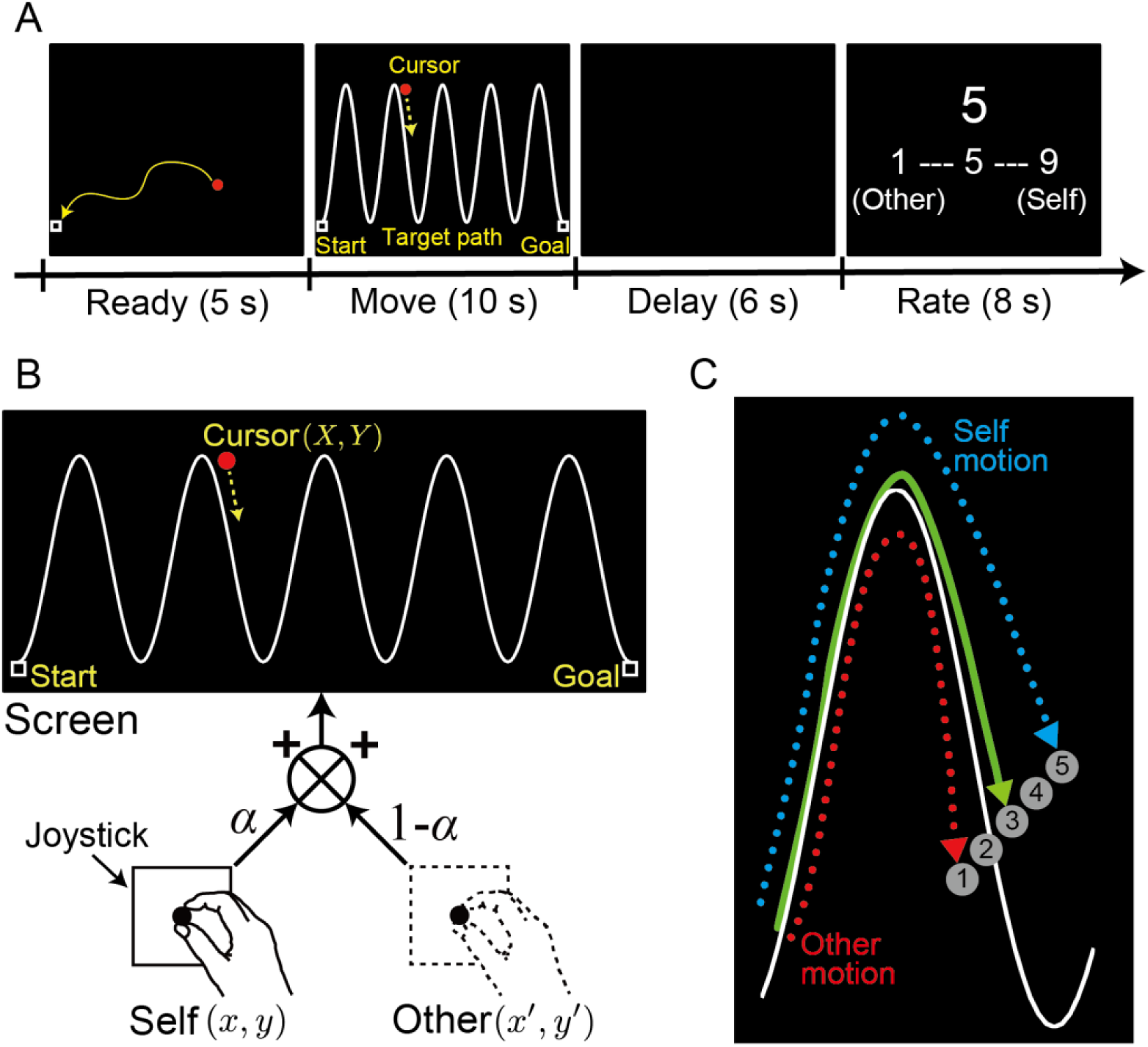
Motion morphing task. (A) Trial timeline. After moving a cursor to the start position (shown as a square) during the 5-s Ready period, participants traced a sinusoidal target-path with a cursor controlled by a joystick during the 10-s Move period. Following a 6-s Delay period, participants selected a rating score (a 9-point Likert scale) about their self-other attribution by pushing buttons. (B) Morphing method. Cursor position on the screen (*X*, *Y*) was the weighted summation of the joystick position (*x*, *y*), which is controlled by the current participant (self), and pre-recorded joystick position (others) (*x*’, *y*’). The weights were changed by a morphing ratio (α). (C) Cursor trajectories. The circles labeled with numbers (1–5) illustrate how the cursor position was changed according to the five morphing ratios (0 ≤ *α* ≤ 1): self 0% (number 1) to self 100% (number 5) with every 25% step. In the self-other mixed conditions, the cursor was displayed (number 2, 3 or 4) between the position of their own joystick and the position of the other person’s joystick.

### Relationship between tracing behavior and a rating score of self-other attribution

We first identified the sensorimotor factors, which affect the participants’ rating score on the self-other attribution. We investigated the relationship between trial-by-trial rating scores and four individual behavioral measures: (1) the target-cursor error, which is the vertical distance between the target path and the cursor position (the blue arrow in the left panel in Fig. 3A), (2) the target-joystick error, which is the vertical distance between the target path and the joystick position (the green arrow in the left panel in Fig. 3A), (3) the cursor-joystick positional error, which is the Euclidean distance between the cursor and the joystick position (the red arrow in the left panel in Fig. 3A), and (4) the cursor-joystick velocity error, which is the absolute difference in velocity between the cursor and the joystick movement (the orange arrow in the right panel in Fig. 3A). We calculated the Fisher-transformed Pearson’s correlation coefficients between each behavioral measure (mean value within every second) and self-other rating scores (one value for each trial). Figure 3B shows a time course of the correlation for self 50% condition (Supplementary Fig. S2 shows time courses for all conditions). In the self 50% condition, the upper limit of 95% confidence interval (CI) of the cursor-joystick positional error was lower than zero from 2 s after the onset of the movement (red line in Fig. 3B). Similarly, the 95% CI of the velocity error was less than zero from 3 s after the onset of the movement (orange line in Fig. 3B). This negative correlation indicates that the more the cursor- joystick positional or velocity error was, the more likely were the participants to judge the cursor movements to be attributed to other’s motion, and vice versa. The negative correlation gradually became larger according to error accumulation, which was calculated as the mean from movement onset to every second (red and orange lines in Fig. 3C show for self 50% condition, and in Supplementary Fig. S3, show for all conditions). This result indicates that the accumulation of error between cursor and joystick is important for the judgment of self-other attribution. By contrast, the correlation coefficients for the target-cursor and target-joystick errors were stable around zero (blue and green lines in Fig. 3B and 3C).

**Figure 3.**
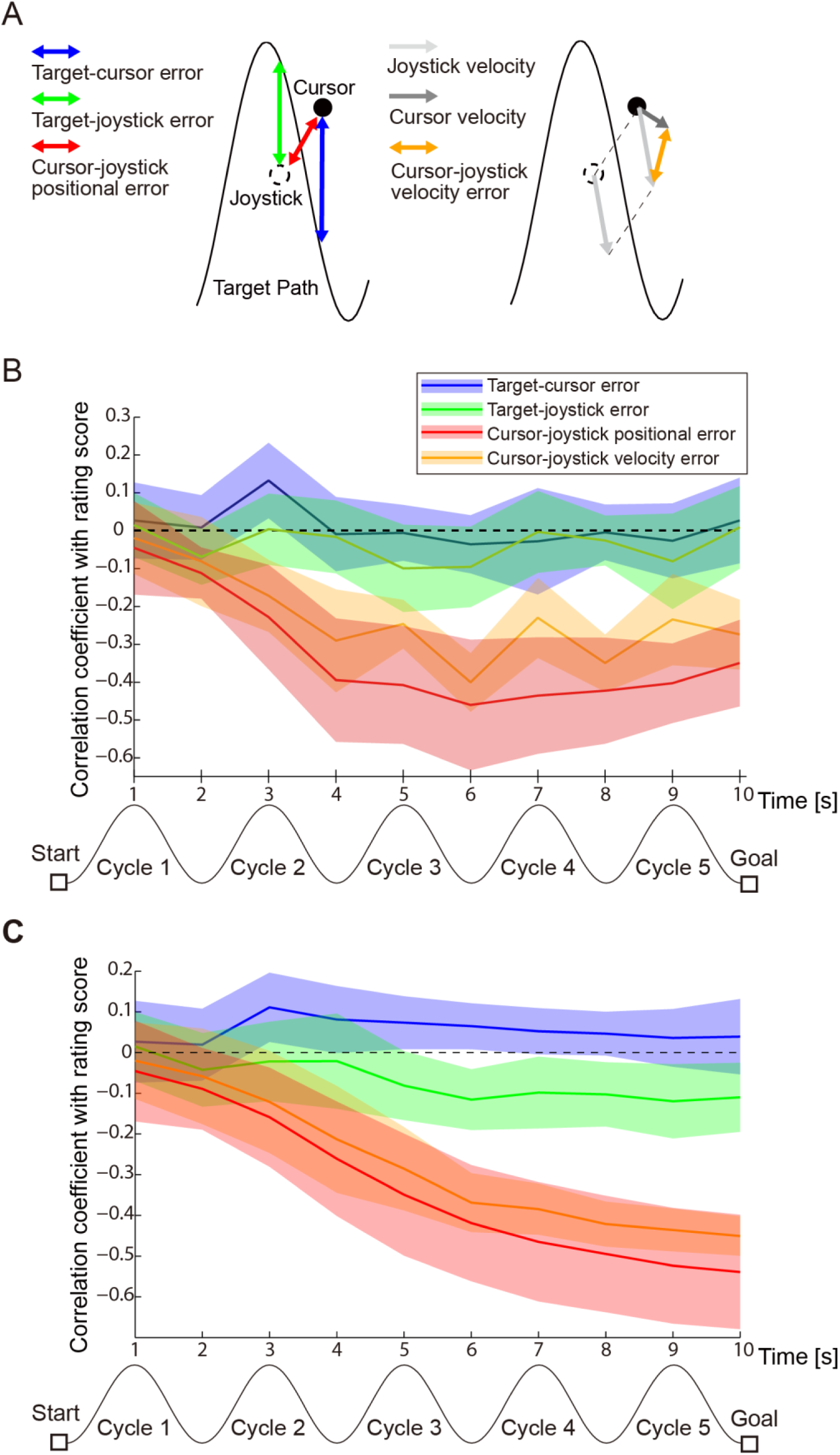
Relationship between tracing behavior and a rating score of self-other attribution of movement. (A) Schematic of the four behavioral measures whose relationship with the self-other attribution score was investigated. The blue, green and red double-headed arrows in the left panel indicate the target-cursor error, the target-joystick error, and the cursor-joystick positional error, respectively. The light and dark gray arrows in the right panel denote the joystick and the cursor velocity, respectively. The orange double-headed arrow represents the cursor-joystick velocity error. (B) Time courses of Fisher-transformed Pearson’s correlation coefficients between each behavioral measure and self-other rating scores during the 10-s Move period. The values of behavioral measures were averaged within every second. Colored shaded area denotes 95% confidence intervals. Note that the data in self 50% condition is shown. Negative correlation indicates that the more the error was, the lower the score (i.e., more other-attribution) the participants selected. (C) Time courses of Fisher-transformed Pearson’s correlation coefficients between accumulated value of each behavioral measure and rating scores. The values of behavioral measures were averaged from movement onset to every second. Colored shaded area denotes 95% confidence intervals. Note that the data in self 50% condition is shown. Negative correlation indicates that the more the error was accumulated, the lower the score the participants selected.

Although the joystick position was not displayed on the screen, participants could predict the actual position and velocity of their joystick on the screen according to their proprioception and a forward model of the relationship between the cursor and joystick acquired during the practice run under the self 100% condition (see Methods: Behavioral task for more details). A cursor appeared in the position shifted by addition of the other’s joystick position to the predicted position (i.e., actual position) in the self 0 – 75% of conditions (Fig. 2C). Therefore, the cursor-joystick positional and velocity errors were reasonably regarded as proxies for a prediction error of sensory feedback. Time courses of correlations between the errors and the self-other rating score (red and orange lines in Fig. 3B and 3C) indicate that the accumulation of prediction error explains a large part of variance in self-other attribution.

### Decoding “prediction error”

Having identified the behavioral measures, which are proxies for a prediction error, we investigated the neural correlates of a prediction error. We decoded the cursor-joystick positional and velocity errors from a regional multi-voxel pattern (searchlight analysis^33^, see Methods: Decoding “prediction error” with a multi-voxel pattern regression for more details). We first trained a linear support vector regression (SVR) decoder with the trials for self 100% and 0% conditions and then applied the trained decoder to the trials in self 75%, 50% and 25% conditions, separately. Notably, we averaged the cursor-joystick error from 3 s to 9 s after the onset of the Move period to exclude periods when cursor position was not clearly visible on the screen (see Methods: Task procedure for the detailed description). We predicted the mean error from the multi-voxel pattern by using SVR. As a result, we could decode the cursor-joystick positional error not only in the sensorimotor area but also in the parietal and visual regions (the magenta regions in Fig. 4A, *p* < 0.01 FWE-corrected at cluster-level with a cluster forming threshold *p* < 0.0005). Similar results were obtained for the velocity error (the cyan regions in Fig. 4B). In the first cycle, we could not find any clusters showing significant decoding performance. This is reasonable because the cursor was not displayed for the first two seconds. These results indicate that the broader areas of the brain are related to prediction error information.

**Figure 4:**
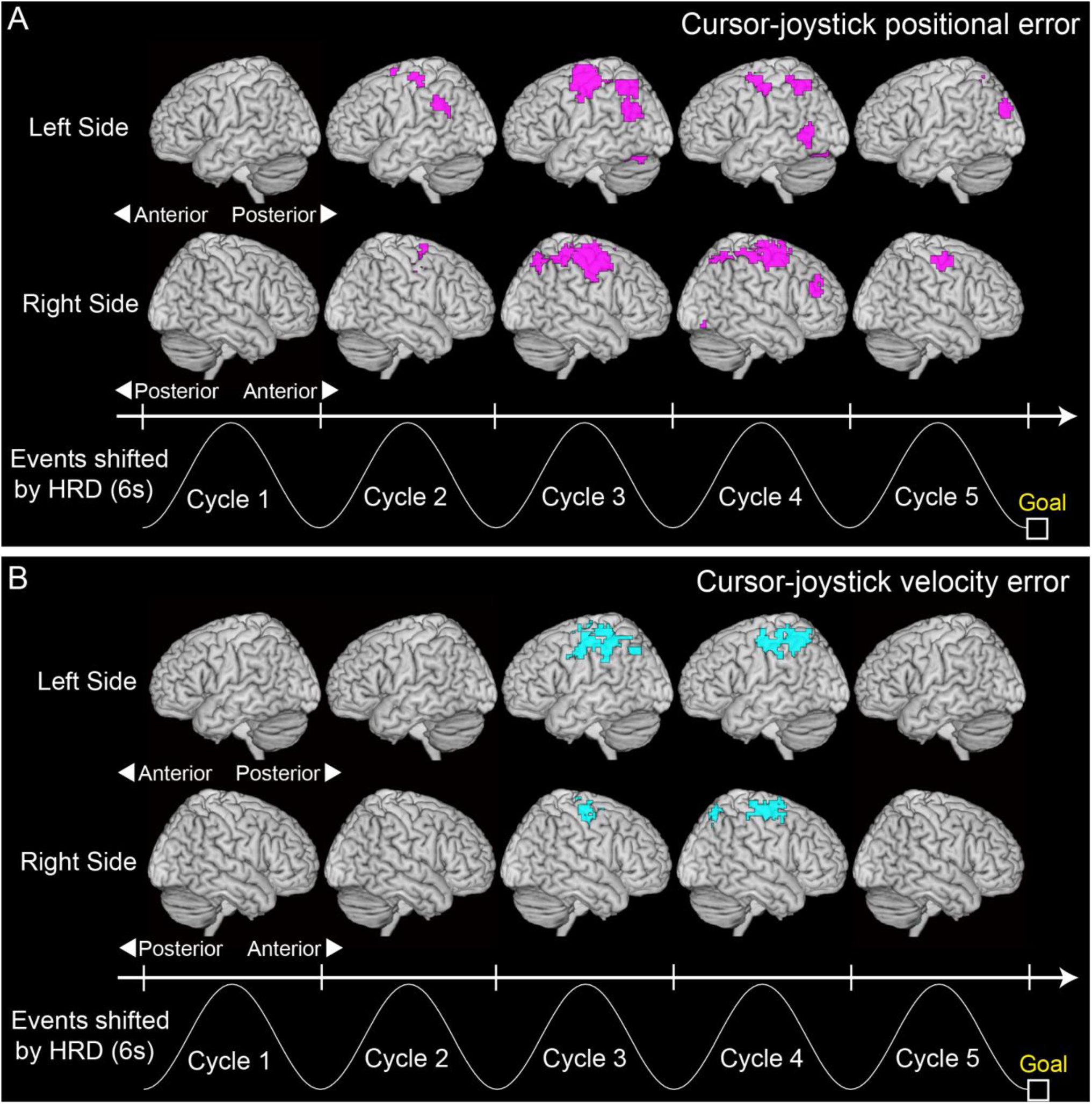
Decoding prediction errors during Move period with multi-voxel pattern regression. (A) Clusters showing significant prediction accuracy for cursor-joystick positional error (magenta regions; *p* < 0.01 FWE-corrected at cluster-level with a cluster forming threshold *p*< 0.0005) and (B) for cursor-joystick velocity error (cyan regions). A searchlight decoding analysis^33^ was applied to a volume data scanned every 2 s during the 10-s Move period to create accuracy maps. The sinusoidal waves represent a typical cursor movement along the timeline shifted by 6 s from the actual time considering the hemodynamic response delay (HRD). All clusters larger than 50 voxels are reported.

### Multi-voxel pattern classification of self-/other-attribution during movement

We decoded self-other attribution of movement, which were evaluated by the participants after movement, from fMRI voxel patterns during the tracing. The participants could not always evaluate with a clear gradation from self to other in the self-other mixed conditions although they were asked to make judgments on a Likert scale. Therefore, we chose classification analysis rather than regression analysis to investigate the predictability of self-/other-attribution. We first allocated the self- or the other-attribution label to the trials with a higher or a lower rating score, respectively, than the mean score across the self 100% and 0% conditions (mean: 5.42, SD: 0.33 across participants). The support vector machine (SVM) classifier was trained with the trials in self 100% and 0% conditions in which distinct activity patterns were expected to appear. Then the classifier was tested for its prediction accuracy with the trials in self 75%, 50% and 25% conditions, separately (see Methods: Decoding self-other attribution with a multi-voxel pattern classification for more details).

A searchlight analysis^33^ found clusters in which self-/other-attribution could be significantly discriminated from their voxel patterns (both white and red regions in Fig. 5A, *p* < 0.01 FWE-corrected at cluster-level with a cluster forming threshold *p* < 0.0005; all the clusters are reported in Supplementary Table S1). At first, the cluster in the basal ganglia (pallidum/putamen regions) appeared at the second cycle in the Move period. Then, in the middle cycles, the right sensorimotor areas including the premotor, primary motor and primary somatosensory cortices (included in pre and post central gyrus) and the left posterior parietal cortices continuously showed significant classification accuracies from the third to the fourth cycles. Only in the last cycle, the right IPL, right medial occipital gyrus (V5/MT+) and supplementary motor area (SMA, included in the posterior medial frontal cortex, see Supplementary Table S1) showed the significant accuracies. These results indicate that not only the posterior parietal regions, which were frequently reported in previous studies^13-15,17,34,35^, but also the sensorimotor and higher visual areas contained information that could predict the following self-/other-attribution, and that the neural representation shifted over time during the tracing.

**Figure 5:**
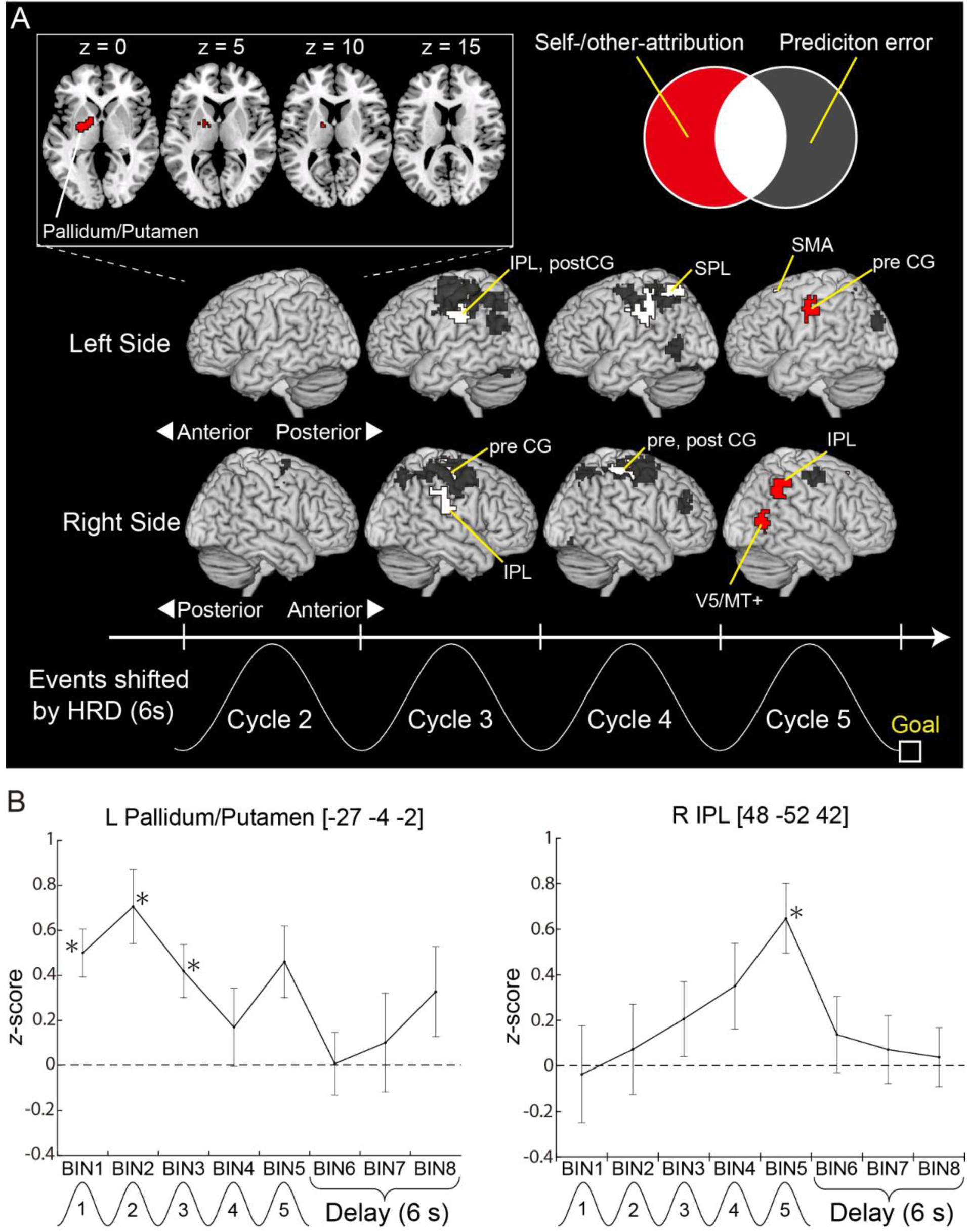
Decoding performance for self-/other-attribution during movement. (A) The clusters showing significant classification accuracy (red and white regions; *p* < 0.01 FWE-corrected at cluster-level with a cluster forming threshold *p* < 0.0005). The gray regions denote areas showing significant prediction accuracy for cursor-joystick positional and velocity error (the gray area of the Venn diagram in the upper panel), which correspond to the colored regions in Fig. 4. The red regions denote areas showing significant accuracy in self-/ other-attribution (the red area of the Venn diagram). The overlapping regions are colored white (the white area of the Venn diagram). The sinusoidal wave represented at the bottom is shifted by 6 s from the actual time considering the HRD. All clusters larger than 50 voxels are reported. SPL: superior parietal lobe, IPL: inferior parietal lobe, pre CG: precentral gyrus, post CG: postcentral gyrus, SMA: supplementary motor area, MT: middle temporal. (B) Time courses of *z*-scores (i.e., classification performance) at the peak voxel in the left pallidum/putamen and right IPL. Each time bin corresponds to a volume data scanned every 2 s during the 10-s Move and 6-s Delay periods. The events denoted under the time bins are shifted by 6 s from the actual time considering the HRD. Asterisks indicate *z*-scores that were significantly larger than zero (*p* < 0.01 uncorrected, two-tailed one-sample *t*-test).

However, it is still unclear whether the classification was based on a prediction error (sensorimotor processing) or self-other rating score (conscious experience of self-agency). We overlapped the regions where we could decode the positional or velocity error (gray clusters, corresponding to the gray area in the Venn diagram in Fig. 5A) on the regions where we could predict self-/other-attribution. We found overlaps in the sensorimotor and superior parietal regions (white clusters, corresponding to the white area in the Venn diagram); however, there exist regions showing significant performance solely in decoding self-/other-attribution. Such regions (red clusters, corresponding to the red area in the Venn diagram) were found in the left pallidum/putamen (at the second cycle), left precentral gyrus, right V5/MT+ and right IPL (at the fifth cycle).

Next, we confirmed, by using a multivariate regression analysis, whether the activity patterns in the above four regions can directly decode the self-other rating score (without the above classification). We found that the decoding performance in the left pallidum/putamen (*t*_10_ = 3.88, *p* = 0.012 Bonferroni corrected, two-tailed one-sample *t*-test for *z*-scores of the correlation coefficients after permutation) and right IPL (*t*_10_ = 3.96, *p* = 0.011 Bonferroni corrected) reached significance although those in the left precentral gyrus and right V5/MT+ did not (*t*_10_ = 2.10, *p* = 0.25 and *t*_10_ = 2.45, *p* = 0.14 Bonferroni corrected, respectively). Taken together, the above findings suggest that neural representation in the left pallidum/putamen and right IPL are more sensitive to conscious experience of self-other agency as compared to prediction error.

We further investigated the temporal changes in classification performance (*z*-score, see *Evaluation of classification accuracy in individual analysis* in Methods section) at the peak voxels of the clusters in the left pallidum/putamen and right IPL (Fig. 5B). In the left pallidum/putamen, the *z*-score reached the peak at around the second cycle (time bin 2) and gradually decreased over time. Meanwhile, performance in the right IPL gradually increased and reached the peak at around the fifth cycle (time bin 5). These time courses indicate that neural information, which could discriminate self-/other-attribution, only appeared during the Move period; however, the timing of the appearance was different between the left pallidum/putamen and right IPL.

### Multi-voxel pattern classification of self-/other-attribution after movement

We applied the classification analysis to the data during the Delay period (the interval between the offset of Move period and the onset of Rate period). As a result, the classification accuracies in the right IFG, specifically Brodmann areas 44 and 45, and VLPFC were significantly above the chance level (Supplementary Fig. S4A, *p* < 0.01 FWE-corrected at cluster-level with a cluster forming threshold *p* < 0.0005; all the clusters are reported in Supplementary Table S2). It might be possible that activity for preparation of an action with their right hands to select a rating score in the Rate period contributes to the high accuracy in the left sensorimotor areas. However, it is remarkable that the significant accuracies in the right prefrontal regions appeared only after the Move period.

## Discussion

The current study explored the neural representation corresponding to individual processes in a theoretical concept of a hierarchical system for a sense of agency (Fig. 1). We first found that attribution judgments were tightly linked to accumulation of a prediction error, which resulted from comparison between predicted and actual sensory feedback (Fig. 3B and 3C). The MVPA revealed that the widespread area of the sensorimotor and parietal regions represented the prediction error information from the early stage of the Move period (Fig. 4). Secondly, in classifying self-/other-attribution, we found the regions in the sensorimotor, posterior parietal and higher visual cortices to contain discriminative information. Most of the regions overlapped those related to prediction error (i.e., low-level sensorimotor information); however, the left putamen and right IPL showed significant performance solely in decoding self-/other-attribution (Fig. 5A). Finally, the voxel patterns in the right IFG and VLPFC could significantly discriminate self- from other-attribution only after movement (Supplementary Fig. S4A). Our findings demonstrate that the brain develops distinct types of information with the passage of time to build up the sense of agency.

The theoretical models^23,24,36^ suggest that a hierarchical system consisting of (1) sensorimotor, (2) non-conceptual feeling and (3) conceptual judgment processes, is fundamental to the sense of agency (Fig. 1). We found that prediction error information and self-other attribution could be predicted by the voxel patterns in the different clusters respectively (Fig. 5A). We assumed that the regions, which can decode prediction error information, can contribute to the first sensorimotor process of the hierarchical system. Meanwhile, those, which can predict self-other attribution, are responsible for higher-order functions, rather than low-level sensorimotor processing. We also found the distinct areas which show significant classification performance in self-/other attribution during the movement (Fig. 5A) or after the movement (Supplementary Fig. S4A). Considering the role of feeling of agency to take over sensorimotor information from the first process, the neural processing for feeling of agency might appear before the processing for the judgment of agency. Therefore, it is plausible that the regions, which could classify self-/other-attribution during movement, mainly contribute to the second non-conceptual feeling process, while those, which could classify self-/other-attribution after movement, are highly involved in the final judgment process.

A comparator model proposed that a prediction error is the main factor determining action attribution^21,23^. Consistent with the model, our data demonstrated the tight relationship between trial-by-trial cursor-joystick positional and velocity errors, which were proxies for a prediction error, and self-other rating score (Fig. 3B and 3C). Such errors could be predicted by voxel patterns in the widespread areas of the sensorimotor, posterior parietal and visual cortices (Fig. 4). Thus, these were mainly responsible for the first sensorimotor process.

We found the two clusters, the left putamen (in the second cycle) and the right IPL (in the last cycle) as regions where we could solely decode self-/other-attribution but not decode prediction error information (Fig. 5A). However, we conclude that the right IPL is only a plausible candidate to reflect feeling of agency based on prediction error in the hierarchical system (Fig. 1) according to the following reason. While the classification performance in the right IPL gradually increased during movement, the performance in the left putamen reached the peak at the second cycle (Fig. 5B). In the second cycle, a correlation between a prediction error (cursor-joystick positional or velocity error) and a self-other rating score was relatively low (Fig. 3B), and information on the prediction error was not sufficiently accumulated to be reflected in a rating score (Fig. 3C). Moreover, the putamen is mainly involved in automatic and unconscious execution of learned movements^37,38^. Due to the repetitive execution in the practice and main runs under the self 100% condition, participants were accustomed to trace the target line smoothly with the normal cursor. Taken together, the activity patterns in the left putamen probably reflect such smoothness of cursor control rather than a feeling of agency based on prediction error.

Previous studies implicate that the right IPL contributes to the process for non-conceptual feeling of agency. First, the right IPL is the region most frequently reported as a neural correlate of a sense of agency in previous neuroimaging studies^13-15,17,34,35^. Second, a single pulse transcranial magnetic stimulation (TMS) over the right IPL induced a change in the participant’s action attribution^39-41^. Finally, Chambon et al.^42^ demonstrated that the right IPL reflects subjective experience of self-agency even without receiving a prediction-error signal calculated in the sensorimotor system^11,39,43,44^. They reported that the prospective signals, particularly the fluency of action selection controlled by subliminal priming, affected the sense of agency^45^. Their fMRI study suggested that a region in the right angular gyrus, which is a part of the right IPL, received the prospective signals carried from the dorsolateral prefrontal cortex and contribute to a subjective experience of self-agency^42^, thus suggesting that the right IPL is responsible for higher-order function, rather than low-level sensorimotor processing, in the system for the sense of agency. In our case, the right IPL received prediction error information processed in the broader areas of the sensorimotor regions to make participants experience self-agency.

We found high classification performance in the right IFG (specifically Brodmann areas 44 and 45) and VLPFC only during the Delay period, which was followed by the Rate period (Supplementary Fig. S4A). Because participants were likely to determine their rating score in this period, we assumed that these regions were mainly responsible for the judgment process. This assumption is supported by the fact that the time courses of classification performance (*z*-score) reached the peak value only after movement at the peak voxels in the right IFG and VLPFC (Supplementary Fig. S4B). In addition, these right inferior frontal cortices are known to be recruited for recognition of self-face or self-bodily features^6,7,9,46-49^. Self-recognition requires one to evaluate bodily features in relation to perceptual or mental image of oneself and to attribute them to self or other^2^. These studies suggest that the right IFG and VLPFC might contribute to the conceptual judgment step in the hierarchical system (Fig. 1).

The clusters we found in the right prefrontal cortices are parts of the right inferior branch of the superior longitudinal fasciculus tract (SLF III)^50^. The IFG is densely connected with the right IPL through the inferior part of the SLF III ^51^. Importantly, this fronto-parietal network contributes not only to recognition of self-body^47-49^ but also to awareness of our bodily sensations^47,52^. These findings proposed the idea that the network is a neural basis of bodily self-consciousness^49^. The right IFG and VLPFC (the third process) densely communicate with the right IPL (the second process) to build conceptual judgment of agency by referring to feeling of agency represented in the right IPL.

There exist several possible confounding factors in the MVPA. The first factor is the difference in attention level to control the cursor whether the participants felt the cursor movement was attributed to self or controlled by other. The possible scenario would be as follows: participants might find it more difficult to precisely control the cursor in the presence of an external agent controlling the cursor. This would suggest that the more strongly the participants felt the cursor was controlled by an external agent, the more attention they might have paid towards controlling the cursor. In that case, we could have only decoded different levels of attention covarying with action attribution. We checked if the attention level correlated with a rating score of the self-/other-attribution. Although we cannot directly measure the level of attention on cursor control, it can be inferred from accuracy in participant’s tracing performance, which was measured by error between the target path and the cursor position (target-cursor error). Note that participants were instructed to precisely trace the target path with the cursor even in self-other mixed condition. The target-cursor error did not correlate with the rating score (the blue line in Fig. 3B), suggesting that the attention on cursor control was not a crucial factor to classify the self- and the other-attribution judgment. The second factor is difference in rating number participants prepared in their mind prior to the Rate period. We instructed them to judge action attribution on a 9-point Likert scale. Although the Rate period was temporally apart from the Move and Delay period, it might be possible that participants kept the rating number in mind before the Rate period. The right IPL was suggested to be involved in a magnitude system of numerical processing^53^. Thus, we might have decoded the difference in the rating number from activity patterns in the right IPL even before the Rate period. However, the cluster in the right IPL disappeared once entering the Delay period (Fig. 5B), suggesting that numerical processing was not the crucial factor of our successful classification.

To answer how the brain organizes multistep processes in a hierarchical system for a sense of agency, we employed the two strategies. The first was the unique experimental paradigm considering temporal evolution of self-agency^25-27^. Most previous studies required participants to perform an intermittent action such as a button press (e.g., reference^15^) or a reaching movement (e.g., reference^12^). However, such simple tasks rather made it difficult to dissociate the multistep processes and, consequently, to clarify a role of different brain regions. In our experiment, participants continuously received sensorimotor evidence while tracing a target path under the ambiguous condition so that they could gradually realize whether cursor movement was attributed to self or other. Thus, our task paradigm enabled us to separately investigate individual processes from sensorimotor processing to cognitive judgment of agency.

The other strategy was to apply a MVPA to fMRI data^28-30^. Almost all the previous studies used the mass univariate rather than multivariate analysis to identify the regions responsible for either self- or other-agency (reviews^18,19^). However, it has been unclear how the brain integrates information represented in different regions, each of which is related to self- or other-agency, and constructs a unified judgment about action attribution. Our MVPA results demonstrate that the right IPL and the right inferior frontal cortices respectively modulated their neural activity patterns to reflect feeling and judgment of self-/other-agency (Fig. 5 and Supplementary Fig. S4). This suggests that these areas act as prime candidates for integrating individual information about self- or other-agency processed in multiple brain regions. We have confirmed that it is unlikely that the MVPA result (Fig. 5) would only reflect the difference in the regional activation level (for more details see Supplementary Results: Mass univariate analysis of voxel-wise activation with self- and other-attribution judgment and Supplementary Fig. S5).

In 1890, William James proposed the concept of the ‘I’ as one aspect of the self: experiencing oneself as a subjective agent of thought, perception and action^1^. Our study has revealed the mechanism underlying awareness of ourselves as an agent of an action in the brain through interaction with the external world, which is, as it were, the neural basis of the ‘I’. Wegner^54^ suspected that self-agency is a trick of the mind induced by a cognitive inference about relationships between thoughts and actions. Our study does not deny the importance of the cognitive inference in the determination of self-agency but demonstrated that results of sensorimotor processing evolve subjective feeling and contributes to the cognitive judgment of agency in the brain. Thus, a sense of agency is not limited to a cognitive system but is grounded in subjective experience based on the sensorimotor system.

## Methods

### Participants

Eighteen right-handed and healthy volunteers with a mean age of 25.9 (20-42 years of age) participated in our experiment (six females). A power analysis for main effect of morphing ratio on the rating score (Supplementary Results: Behavior results and Supplementary Fig. S1) was conducted with power selected at 0.8, effect size (f) at 0.4 and alpha at 0.05. The sample size was chosen as 18 because the analysis determined 16 to be optimum. We eliminated data from seven participants from the analysis to ensure homogeneity of action attribution process (for more details see Supplementary Results: Participants’ rejection from fMRI analysis). Note that the statistical power for the main effect was still high (more than 0.99) even after the exclusion of the participants. We eventually analyzed the fMRI data of eleven participants with a mean age of 25.5 (two females). Written informed consent was obtained from all the volunteers in accordance with the latest version of Declaration of Helsinki. The experimental protocol was approved by the ethics committee at Kochi University of Technology.

### Behavioral task

#### Trial Timeline

Participants were required to trace a five-cycle sinusoidal wave (a target path) with a cursor (Figure. 2A)^26^. Participants manipulated a joystick with their right hand and controlled the cursor on a screen. At the beginning of each trial, the trial number was displayed on the screen for 1 s. During five countdown sounds, they set their cursor at the starting point located near the lower left corner of the screen. After hearing the count “zero” sound, they started to trace the target path toward the goal point located near the lower right corner of the screen, as smoothly and as precisely as possible. They were instructed to trace each half cycle of the wave in 1 s (i.e., 0.5 Hz) and to complete the entire movement within a Move period of 10 s. A blank screen was presented during a Delay period for 6 s. Then participants reported how much they felt the cursor movement to be attributed to their own joystick movement on a 9-point Likert scale from 1 (completely other’s movement) to 9 (completely their own movement). The number 5 was displayed on the screen at the beginning of a Rate period for 8 s (Fig. 2A). The number was incremented or decremented by one with the press of the right or left button, respectively. The buttons were attached to the joystick box, and participants were instructed to press the buttons with their right hand. Note that the cursor was not displayed on the screen during the first 1.5 s and the last 0.5 s of the Move period because the onset and the offset of the cursor movement are sensitive to the mismatch between their own joystick and cursor movement, which predominantly affects self-other attribution judgment.

#### Morphing visual feedback from self to other

Visual feedback (i.e., cursor movement) during the tracing movement was morphed by incorporating other person’s movement into the participant’s movement. We calculated the weighted summation of the participant’s online joystick position (*x*, *y*) and other’s position (*x*’, *y*’) at 60 Hz (the refresh rate of the monitor) and displayed the cursor in the calculated position (*X*, *Y*) on the screen (Fig. 2B). The weight corresponded to the morphing ratio (α). Other persons’ movement was recoded prior to the fMRI experiment, and 240 trajectories (fifteen trajectories recorded from each sixteen participant) were stored in the data pool. A trajectory was randomly chosen for each trial from the pooled data. There were five morphing ratios: self 0% (other 100%, α = 0) to self 100% (other 0%, α = 1.0) with every 25% step. In the self 100% condition, the visible cursor position corresponded to the participant’s joystick position (the cursor labeled with the number 5 in Fig. 2C). By contrast, in the self 0% condition, the visible cursor position was independent of their own joystick position (the cursor labeled with the number 1 in Fig. 2C). In the self-other mixed conditions, the cursor was displayed at a position between the position of their own joystick and that of his/her pre-recorded joystick (the cursor labeled with number 2, 3 or 4 in Fig. 2C). We instructed the participants to trace the target path with the cursor as closely as possible even if the movement of others was strongly felt during the cursor control.

#### Experimental procedure

Before the main fMRI runs, participants performed two types of practice runs inside the fMRI scanner. In the first practice run, the participants were trained to trace the target path with a cursor in accordance with 1-Hz metronomic sounds to get accustomed to the cyclic movement. In this run, the cursor movement precisely reflected their joystick movement (self 100% condition). In the second practice run, they conducted the same task as the main fMRI runs but the number of trials was smaller (10 trials) than that of the main runs (50 trials/run). After the practice runs, participants conducted three main runs/day (150 trials), and total six runs (300 trials) for two days. The participants underwent the five morphing ratios ten times in a random order during each of the main runs.

### MRI data acquisition

A 3 Tesla Magnetom Verio scanner (Siemens, Germany) with a 32-channel head coil was used to acquire T2*-weighted echo-planar images (EPI). In total, 753 volumes were acquired in each run with a gradient echo EPI sequence under the following scanning parameters: repetition time (TR), 2000 ms; echo time (TE), 30 ms; flip angle (FA), 70°; FOV, 192×192 mm; matrix, 64×64; 30 axial slices; and thickness, 4 mm with a 1-mm gap. T2- weighted turbo spin echo images were scanned to acquire high-resolution anatomical images of the same slices used for the EPI (TR, 6000 ms; TE, 58 ms; FA, 160°; FOV, 192×192 mm; matrix, 256×256; 30 axial slices; and thickness, 4 mm with a 1-mm gap). T1-weighted structure images were obtained with 1×1×1 mm resolution with a gradient echo sequence (repetition time, 2250 ms; echo time, 3.06 ms; flip angle, 9°; matrix, 256×256; 192 axial slices; and thickness, 1 mm without gap).

### Preprocessing of fMRI data

The fMRI data were analyzed using SPM8 (Wellcome Trust Centre for Neuroimaging, London, UCL) on MATLAB. We discarded the first three volumes of the functional images in each run to allow for T1 equilibration. The remaining image volumes were temporally realigned to correct for the sequence of slice acquisition and spatially realigned to the first image to adjust for motion-related artifacts. Rigid-body transformations were performed to align the functional images to the structural image for each subject. The images were spatially normalized with the Montreal Neurological Institute (MNI) (Montreal, Quebec, Canada) reference brain and resampled into 3×3×4 mm cuboid voxels. Note that spatial smoothing was not applied to the data. After linear trend removal within each run, we calculated the percent signal change relative to the mean of activity for each run.

### Decoding “prediction error” with a multi-voxel pattern regression

We performed multi-voxel pattern analysis (MVPA) to investigate whether the brain regions contained enough information to predict the prediction error (the positional or velocity error between the cursor and the joystick). A linear support vector regression (SVR) model implemented in LIVSVM (http://www.csie.ntu.edu.tw/~cjlin/libsvm/) was applied to the voxel patterns in each volume with trial-by-trial prediction error as a dependent variable. Note that we used the mean of the prediction error from 3 s to 9 s after the onset of the Move period to exclude periods when the cursor was not clearly visible on the screen (for more detail see above: Behavioral task). The regression analysis was applied to voxel patterns within a 9-mm radius sphere (see below: *Searchlight decoding over the brain*) extracted from each volume of fMRI data scanned every 2 s (TR = 2 s) during the Move period. Each volume corresponded to a cycle of the sinusoidal movement. The model was trained with data of the trials in the self 100% and 0% conditions and then tested with data of the trials in the other three conditions, separately. We evaluated the decoding performance by using a leave-one-run-out cross validation procedure separately for each participant and each condition. More specifically, we trained the regression model with the fMRI data in five out of six runs and tested with the independent data in the remaining run. This procedure was repeated six times to become each run as test data once.

#### Evaluation of decoding accuracy in individual analysis

We evaluated the above decoding accuracy by calculating *z*-scores of Fisher-transformed Pearson’s correlation coefficient following the permutation procedure^55,56^as follows. We generated 1000-surrogate correlation coefficients by permutating the relationship between the actual and predicted “prediction error” 1000 times to get empirical distribution of correlation coefficient. We calculated the *z*-scores of the original (without the permutation) value based on the empirical distribution. We applied the above steps to each condition of test dataset (self 75%, 50% or 25% condition). We regarded *z*-scores averaged across conditions as decoding performance for each participant before performing a group-level analysis (see below).

#### Searchlight decoding over the brain

We performed a volume-based searchlight decoding analysis^33,57^. We repeatedly extracted voxel patterns within a 9-mm radius sphere containing at least 65 voxels to perform classification analysis. This sphere was moved over the gray matter of the whole brain and was assigned the mean of *z*-scores to the central voxel of a sphere, resulting in the 3-D accuracy map for each participant. The *z*-score maps were smoothed with a 4-mm full-width at half-maximum (FWHM) Gaussian kernel. A random-effects group analysis was performed on the smoothed accuracy maps by using SPM8. We applied a statistical threshold of *p* < 0.01 (FWE-corrected at cluster-level with a cluster forming threshold *p* < 0.0005).

### Decoding self-other attribution with a multi-voxel pattern classification

We performed multi-voxel pattern classification to discriminate the fMRI activity-patterns in trails in which participants attributed the cursor movement to the self from those in which they attributed the movement to the other (see below for label allocation: *Training phase*). We used a binary classifier based on a linear support vector machine (SVM) implemented in LIBSVM with default parameters (a fixed regularization parameter C = 1)^58^. We applied a volume-based searchlight analysis to each volume of fMRI data scanned every 2 s during the Move period.

#### Training phase

We assumed that the typical activity-pattern for judgment on self- or other-attribution appeared in the conditions of an extreme self-other morphing ratio (i.e., self 100% and 0%). Therefore, we trained the classifier using the data in the self 100% and 0% trials. For the binary classification, we defined the border in the rating scores to determine whether participants attributed the cursor-movement to self or other in each trial. We defined the border as the mean of the rating scores across the self 100% and 0% conditions (vertical dashed lines in Supplementary Fig. S6). We labeled trials in the self 100% condition as ‘self-attribution’ if the score was higher than the border (blue bars in Supplementary Fig. S6). By contrast, we labeled trials in the self 0% condition as ‘other-attribution’ if the score was lower than the border (red bars in Supplementary Fig. S6). To equalize the number of trials between the two labels, we selected the same number of trails from each label (down-sampling). Specifically, when the number of ‘self-attribution’ labels was larger than that of ‘other-attribution’ label, the trials with ‘self-attribution’ label were selected in descending order of a rating score to match the number of the trials with ‘other-attribution’ label, and vice versa. We used the data in the selected trials for the training of an SVM classifier.

#### Test phase

For the test phase, the trained classifier was applied to fMRI data in self 75%, 50% and 25% conditions, separately. We used the same criteria as the training dataset to allocate the ‘self-attribution’ or ‘other-attribution’ label to each trial in these conditions. Following the decoding method for “prediction error” (see above), we evaluated the classification performance by using a leave-one-run-out cross validation procedure separately for each participant and each condition.

#### Evaluation of classification accuracy in individual analysis

We evaluated the above classification accuracy in the test phase taking into account the bias due to unequal numbers of trials for the self- and the other-attribution labels in each condition of test dataset (note that the number of trails was equalized between labels in the training dataset, see above). Specifically, we calculated *z*-scores of classification accuracies following the permutation procedure^55,56^as follows. We (1) generated 100-surrogate datasets by permutating the relationship between labels and fMRI data in the test datasets 100 times, (2) applied the trained classifier (see above) to 100-surrogate data to get 100-classification accuracies, (3) randomly sampled 1,000 data from the classification-accuracy dataset allowing duplication to get empirical distribution of classification accuracy, and (4) calculated *z*-scores of the original (without the permutation) accuracies based on the empirical distribution. We evaluated classification accuracies by applying the above four steps to each condition of test dataset (self 75%, 50% or 25% condition). We regarded *z*-scores averaged across conditions as classification performance for each participant before performing a group-level analysis. The resultant 3D accuracy map for each volume scanned during the Move period was smoothed with a 4-mm FWHM Gaussian kernel. A random-effects group analysis was performed on the smoothed accuracy maps. We applied a statistical threshold of *p* < 0.01 (FWE-corrected at cluster-level with a cluster forming threshold *p* < 0.0005).

### Decoding rating score with a multi-voxel pattern regression

We additionally predicted a self-other rating score (ranging from 1 to 9) without classification of the score into the self or other category. The prediction was made from the activation patterns in the four clusters, which showed a significant classification performance of judgment on the self-other attribution but did not show the significant prediction accuracy of either the cursor-joystick positional or velocity error (the left pallidum/putamen, left precentral gyrus, the right IPL and the right V5/MT+; red regions in Fig. 5). We calculated *z*-scores of correlation coefficients between the predicted and actual scores by following the permutation process that evaluated decoding performance in “prediction error”. We performed two-tailed one-sample *t*-test to the *z*-scores across participants to test statistical significance ofthe prediction accuracy.

## Supporting information

## Acknowledgements

R.O. was supported by JSPS KAKENHI Grant Number 15J05135, Grant-in-Aid for JSPS Fellows. R.O. and H.I. were supported by JSPS KAKENHI Grant Numbers 26120002 and 18H01098. The authors are grateful to Dr. Nobuhiro Hagura (The Center for Information and Neural Networks) and Dr. Shu Imaizumi (The University of Tokyo) for their insightful comments.

## Author contribution statement

R.O., T.A. and H.I. designed the study; R.O., T.A., H.K. and H.S. collected its data; R.O., T.A., K.O. and H.I. analyzed the data; R.O., T.A., and H.I. wrote the manuscript; H.K., H.S. and K.O. reviewed and approved the final version of the manuscript.

## Additional information

Supplementary Information

Competing financial interests: The authors declare no competing financial interests.

