## Supplementary material for "Self-agency built with sensorimotor processing: Decoding self-other action attribution in the human brain"

### Supplementary Results

#### Behavior results

Supplementary Figure 1 shows the rating score averaged across all participants as a function of morphing ratio. We found a significant main effect of morphing ratio ($F$ (4, 50) = 152.4, $p < 0.001$) according to a one-way ANOVA with the five morphing ratios as a within-subject factor. The rating scores linearly increased as the ratio of self-movement increased, which was also found in a previous study[1]. Asai[1] fitted the linear regression model to each participant's rating scores and calculated a regression slope to measure individual's self-other discriminability. We similarly fitted a linear regression model to our data; (rating score) = $w_0 + w_1 \times$(self-movement ratio). Note that $w_0$ and $w_1$ indicate a bias parameter and a slope, respectively, and that five self-movement ratios were converted into integer values ranging from zero to four. The mean of the slope was 5.63 (SD: 1.22), which was significantly larger than zero (two-tailed $t$-test: $t(10) = 15.4$, $p < 0.001$).

#### Participants' rejection from fMRI analysis

Asai[1] found that a low slope (low self-other discriminability) is associated with positive schizotypal symptomatology. Therefore, we excluded participants with a significantly low and unstable discriminability from the following fMRI data analysis to ensure homogeneity of action attribution process across participants. In order to systematically determine our exclusion criteria, we investigated an empirical distribution of the slope from an independent dataset including 42 participants. These participants were not enrolled in the current study but conducted the same behavioral task as that in the current study (unpublished data). As a result, the lower limit of the 99.99% confidence interval (CI) of the slope was 3.89 in the 42-participant data. We excluded four participants whose slope was lower than 3.89 from the current fMRI data analysis (red dots in Supplementary Fig. S7A). In addition, we checked stability of our participants' rating score in the most apparent other-attribution (self 0%) condition. We excluded three participants (in addition to the above four) because their SDs of the rating score were higher than the upper limit of the 99.99% CI (2.89) calculated from the 42-participant data (red dots in Supplementary Fig. S7B). Supplementary Table S4 denotes the slope and the SD values of each participant.

**Mass univariate analysis of voxel-wise activation with self- and other-attribution judgment**

We conducted the conventional mass univariate analysis to examine whether magnitude of activity reflected the difference between judgements of the self- and the other-attribution. We used SPM8 for the analysis. For individual analysis, we subtracted voxel-wise activation in the trials with the self-attribution label from activation in the trials with the other-attribution label in the training dataset, and vice versa. Note that the training dataset consisted of the trials in self 0% and 100% conditions. Resultant contrast images were smoothed with a 4-mm full-width at half-maximum (FWHM) Gaussian kernel and entered into a random-effects group analysis.

We found the central part of the cerebellum (VIIIA) as the only area that showed the higher activation in trials with the self-attribution label compared with trials with the other-attribution label (the blue areas in Supplementary Fig. S5, $p < 0.05$ FWE-corrected at cluster-level with a cluster forming threshold $p < 0.001$; all clusters are also reported in Supplementary Table S3). By contrast, the right inferior parietal lobe (IPL), the right inferior frontal gyrus (IFG) and the left supplementary motor area (SMA) showed the higher activation in trials with the other-attribution label compared with trials with the self-attribution label (the red areas in Supplementary Fig. S5). The right IPL region overlapped with the temporo-parietal junction, which has been frequently reported in previous studies about attribution of an external caused action (e.g., reference[2]). However, the above regions found by the mass-univariate analysis did not overlap with the clusters identified by the multi-voxel pattern classification of judgment of self-/other-attribution (Fig. 5A). This univariate result indicates that the multi-voxel pattern analysis (MVPA) result was not just the reflection of the difference in the regional activation.

**Supplementary Figures**

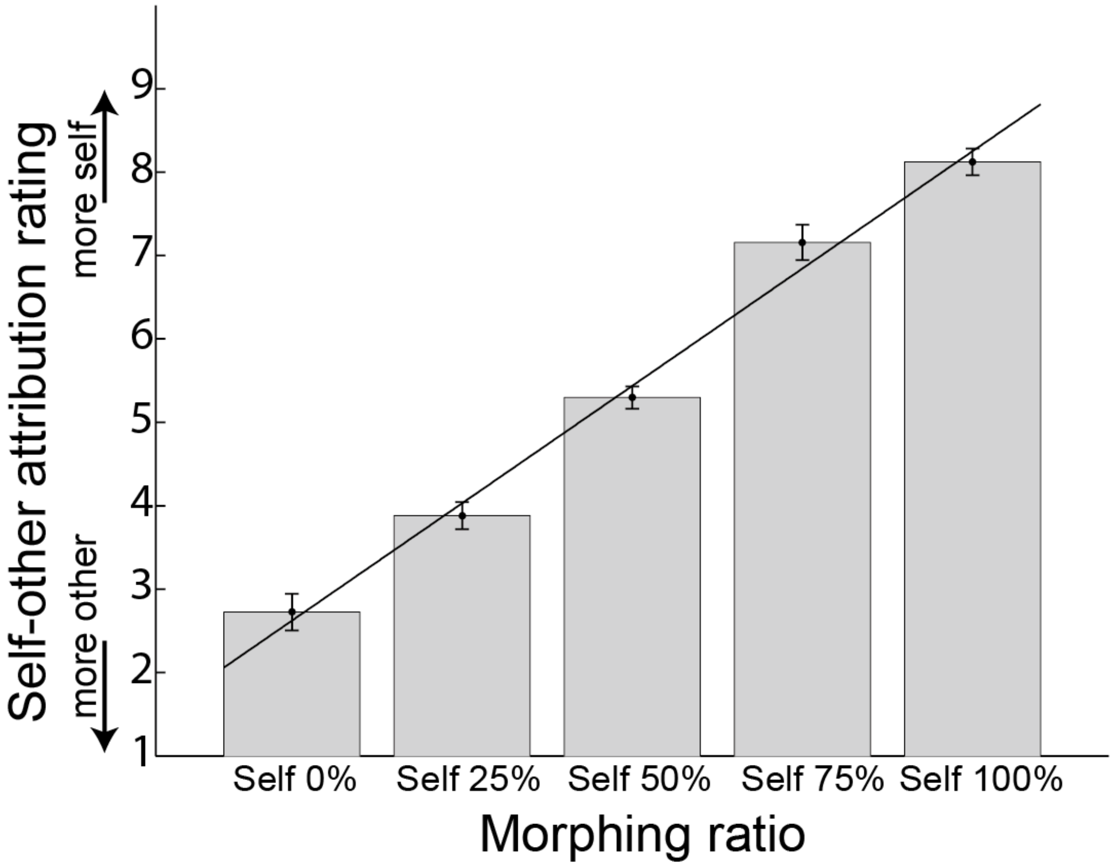


**Supplementary Figure S1. Self-other rating score.**

Self-other attribution rating scores were averaged across participants ($n$ = 11) for each morphing ratio. The higher the score was, the more strongly participants felt that the cursor movement was attributed to their own joystick movement. Error bars indicate SEMs.

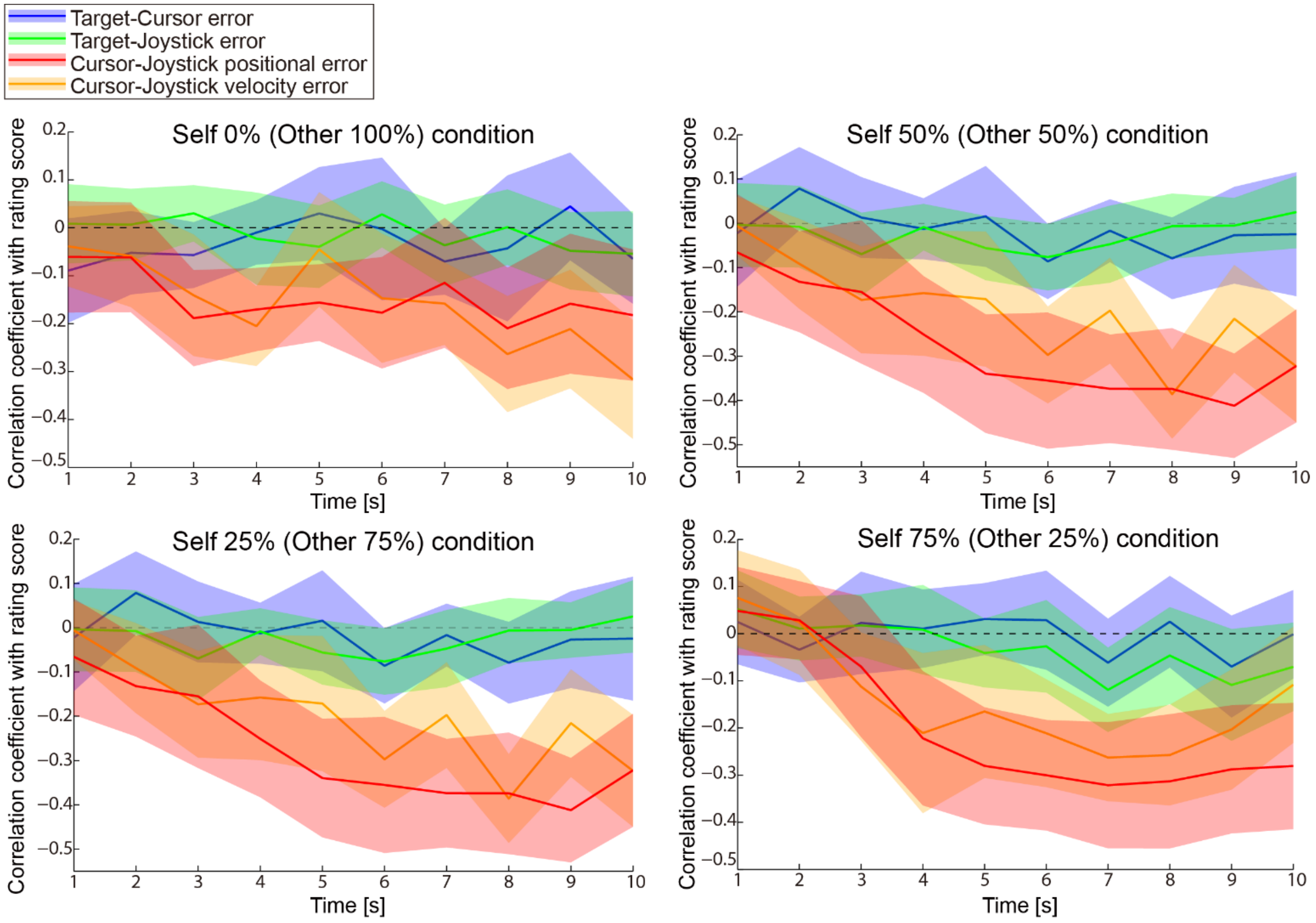


**Supplementary Figure S2: Relationship between tracing behavior and self-other attribution scores (Related to Figure 3B).**

Time courses of Fisher-transformed correlation coefficients between each behavioral measure and self-other attribution scores in all conditions data (except for self 100% condition). Colored lines indicate the positional errors (target-cursor: blue, target-joystick: green, cursor-joystick: red) and the velocity error (cursor-joystick: orange). See Figure 3A for definition of each error. The shaded area denotes 95% confidence interval.

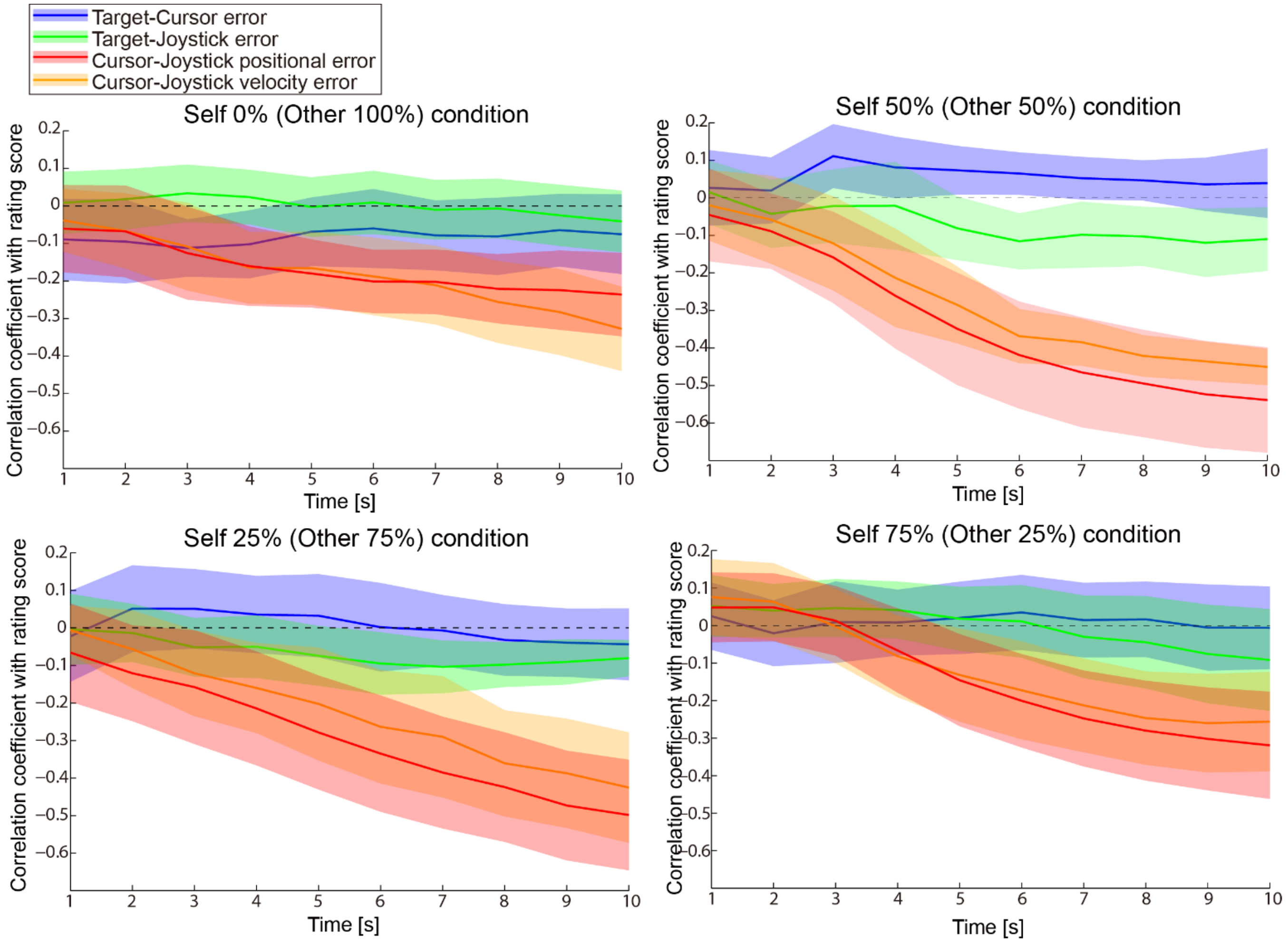


**Supplementary Figure S3: Relationship between accumulated value of tracing behavior and self-other attribution scores (Related to Figure 3C).**

Time courses of Fisher-transformed correlation coefficients between accumulated value of each behavioral measure and self-other attribution scores in all conditions data (except for self 100% condition). Colored lines indicate the positional errors (target-cursor: blue, target-joystick: green, cursor-joystick: red) and the velocity error (cursor-joystick: orange). See Figure 3A for definition of each error. The shaded area denotes 95% confidence interval.

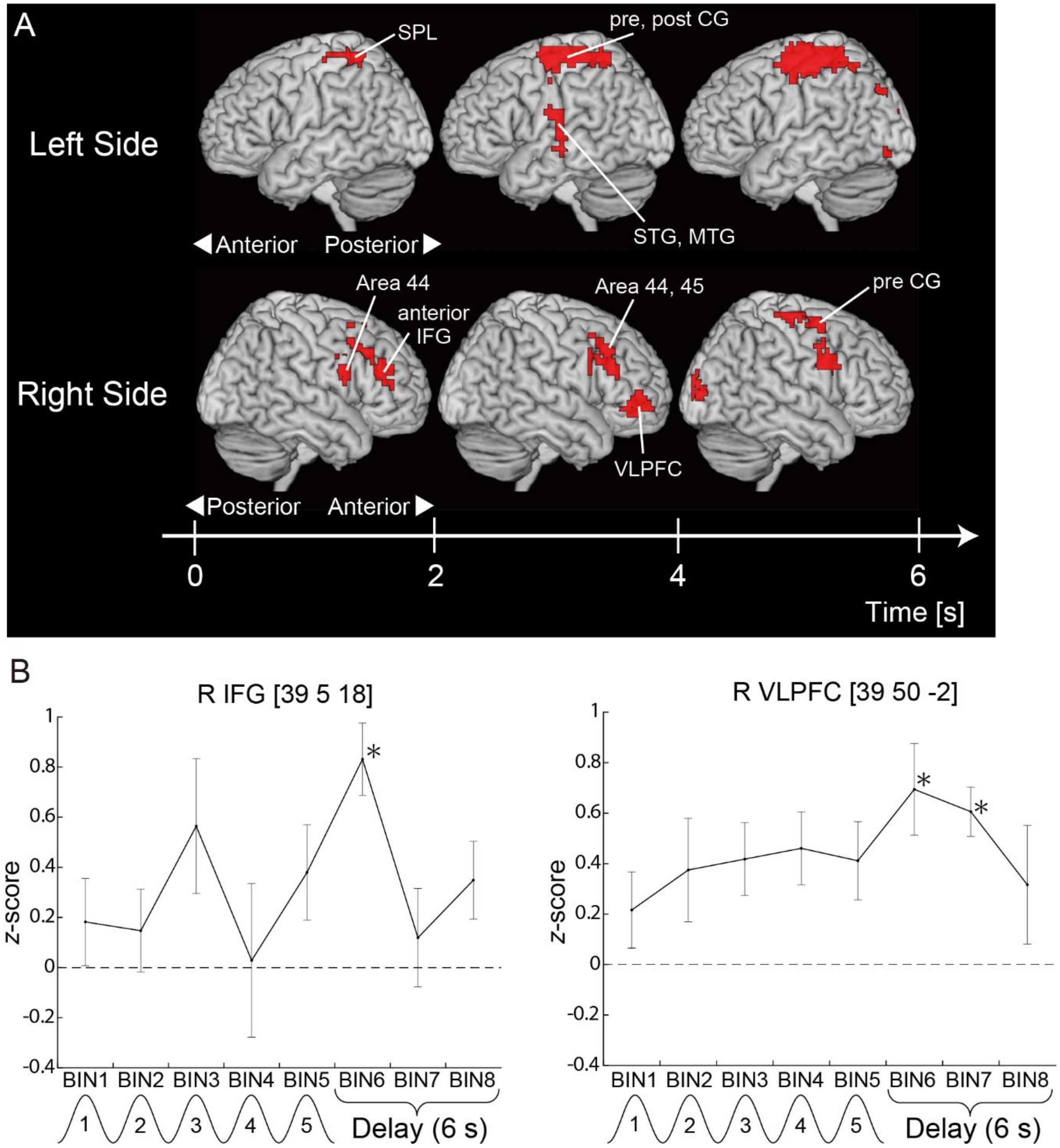


**Supplementary Figure S4. Decoding performance for self-/other-attribution after movement.**

(A) The red clusters show significant classification accuracy ($p < 0.01$ FWE-corrected at cluster-level with a cluster forming threshold $p < 0.0005$). The time axis indicates the time elapsed since the onset of the Delay period. The time is shifted by 6 s from the actual time considering the hemodynamic response delay (HRD). All clusters larger than 50 voxels are reported. IFG: inferior frontal gyrus, MTG: middle temporal gyrus, pre CG: precentral gyrus, post CG: postcentral gyrus, SPL: superior parietal lobe, STG: superior temporal gyrus, VLPFC: ventrolateral prefrontal cortex. (B) Time courses of *z*-scores (i.e., classification performance) at the peak voxel in the right IFG and right VLPFC at around the first 2 s (0-2 s after the Move period) and second 2 s of the Delay period (2-4 s after the Move period), respectively. Each time bin corresponds to a volume data scanned every 2 s during the 10-s Move and 6-s Delay periods. The events denoted under the time bins are shifted by 6 s from the actual time considering the HRD. Asterisks indicate *z*-scores that were significantly larger than zero ($p < 0.01$ uncorrected, two-tailed one-sample *t*-test).

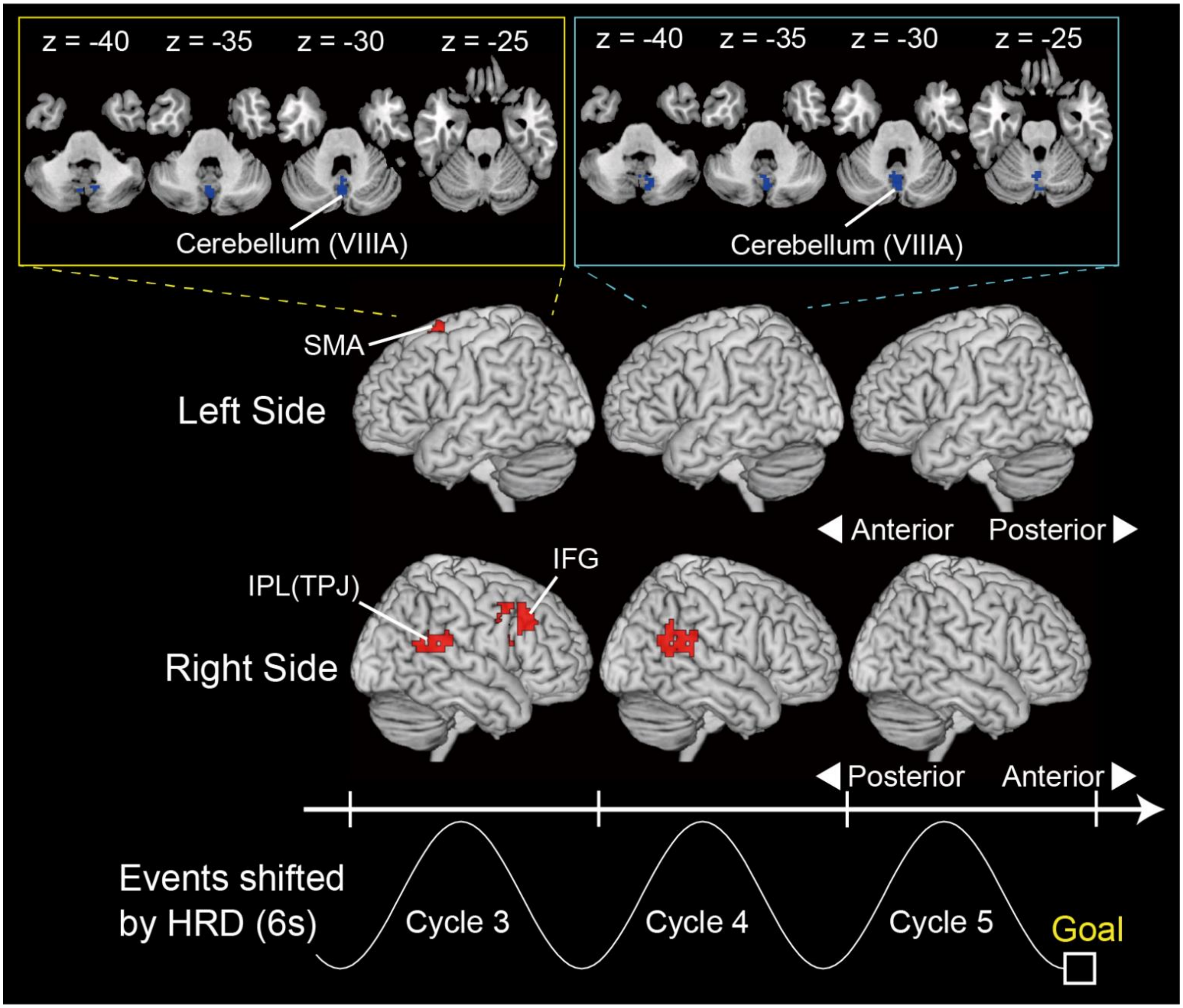


**Supplementary Figure S5. Mass-univariate analysis result during the Movement period.**

The blue clusters (see top panels) showed significantly higher activation when the cursor movement was attributed to self in comparison to when it was attributed to other ($p < 0.05$ FWE-corrected at cluster-level with a cluster forming threshold $p < 0.001$). By contrast, the red clusters showed higher activation when the cursor movement was attributed to other in comparison to when it was attributed to self at the same threshold. Note that the three-cycle sinusoidal wave in the lower panel is shifted by 6 s from the actual time considering the HRD. IFG: inferior frontal gyrus, IPL: inferior parietal lobe, SMA: supplementary motor area, TPJ: temporo-parietal junction.

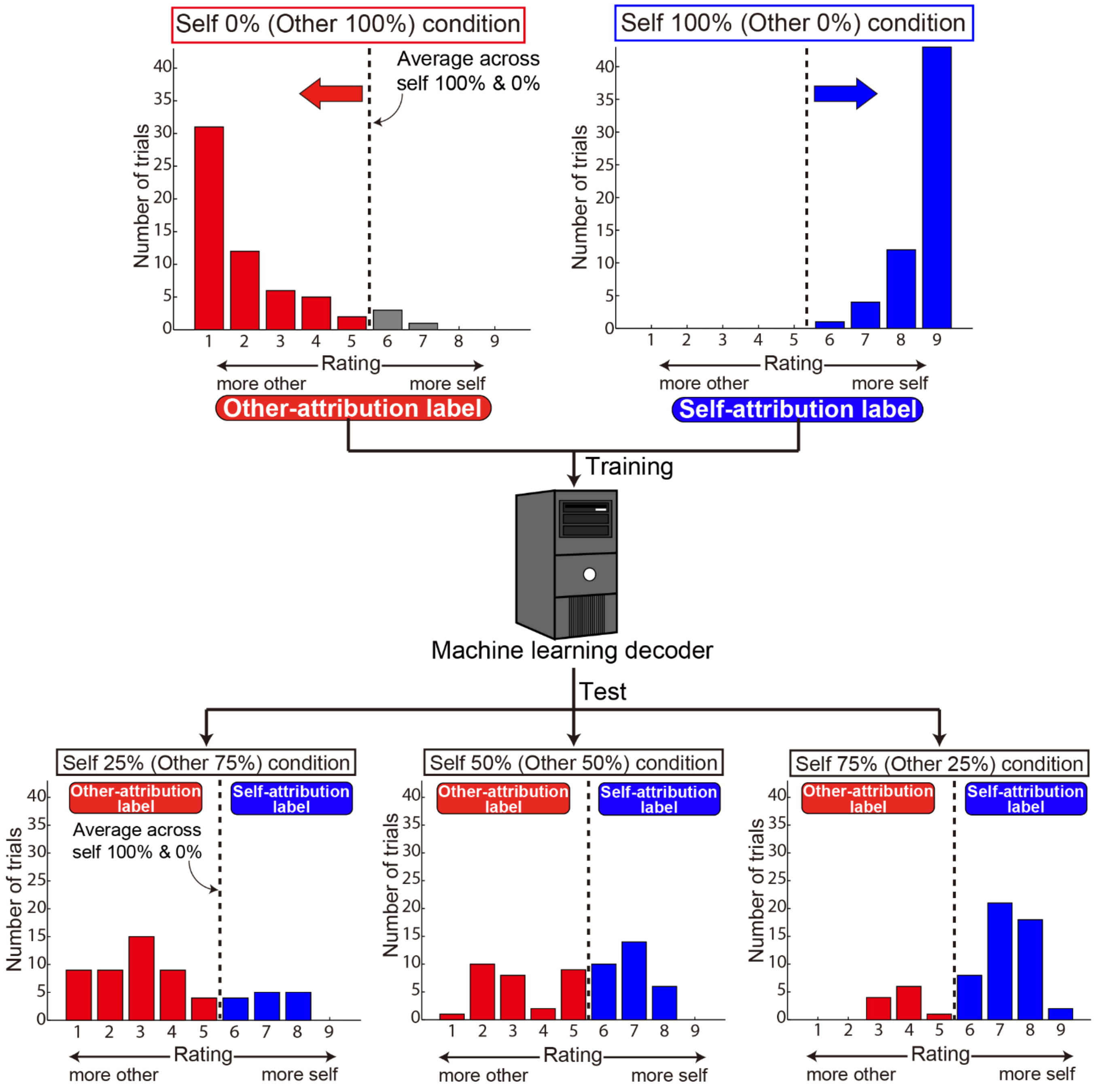


**Supplementary Figure S6. Multi-voxel pattern analysis procedure**

Schematic illustrating the classification analysis procedure. First, the border between self- and other-attribution labels was defined as the average of the rating scores across the trials in self 100% and

0% conditions (the vertical dashed lines). Next, trials in the self 100% and self 0% conditions were labeled as ‘self-attribution’ (blue bars) and ‘other-attribution’ (red bars), respectively, based on the defined border. After down-sampling to equalize the number of trials between the two labels, a machine learning decoder (support vector machine: SVM) was trained. Finally, the trained decoder was applied to fMRI data in self 75%, 50% and 25% conditions, separately, to evaluate classification performance. True labels for test dataset were also allocated using the same border as for the training dataset. Note that the distributions in this figure were acquired from all trials of a single participant.

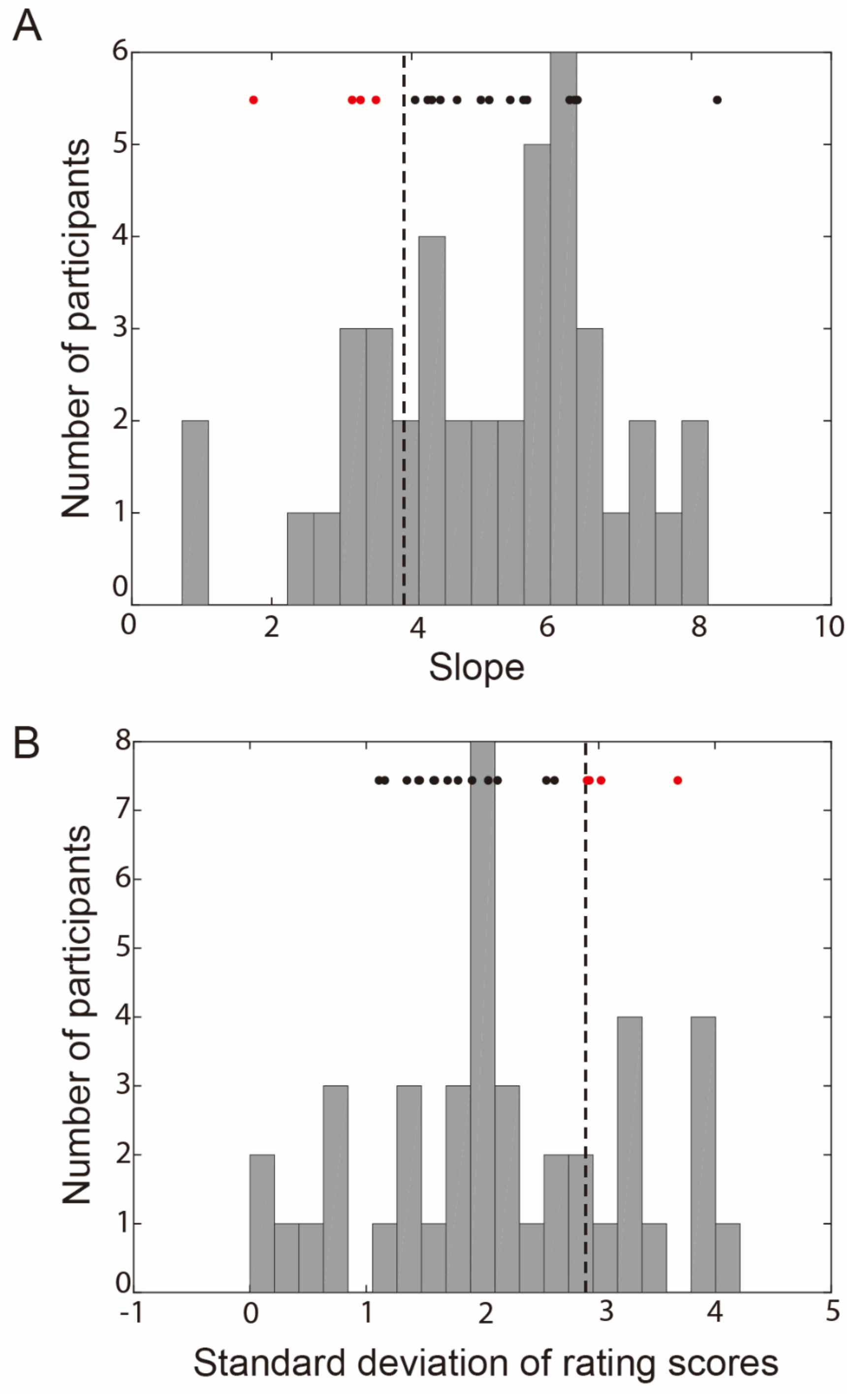


**Supplementary Figure S7. Criteria of participant exclusion based on data from 42 participants**

Bars indicate distribution of slope value of the regression line (A) and standard deviation of rating scores in self 0% condition (B) in 42 participants' data. Dots indicate data from 18 participants participated in the current study. The red dots indicate participants excluded from the main analysis. The dashed vertical line denotes 99.99% confidence interval of each value.

**Supplementary Tables**

| | | | MNI coordinates (peak voxel) | | |
|---|---|---|---|---|---|
| Brain region | Side | Cluster size | x | y | z |
| Cycle 2 | | | | | |
| Pallidum/Putamen | Left | 55 | -27 | -4 | -2 |
| Cycle 3 | | | | | |
| Inferior parietal lobe | Left | 83 | -57 | -25 | 34 |
| Precentral gyrus | Right | 158 | 57 | -22 | 22 |
| Cycle 4 | | | | | |
| Superior parietal lobe | Left | 76 | -27 | -55 | 62 |
| Inferior parietal lobe | Left | 79 | -60 | -28 | 38 |
| Precentral gyrus | Right | 84 | 36 | -25 | 58 |
| Cycle 5 | | | | | |
| Posterior medial frontal cortex | Bilateral | 197 | -9 | -1 | 62 |
| Precentral gyrus | Left | 55 | -45 | -13 | 46 |
| V5/MT+ | Right | 65 | 42 | -58 | 2 |
| Inferior parietal lobe | Right | 52 | 48 | -52 | 42 |

**Supplementary Table S1. Summary of searchlight decoding self-/other-attribution of movement during the Move period**

A threshold at $p < 0.01$ (FWE-corrected at cluster-level with a cluster forming threshold $p < 0.0005$) was set for statistical testing. Clusters larger than 50 voxels are reported. Cycles correspond to those illustrated at the bottom of Figure 5. They are shifted by 6 s from the actual time considering the HRD.

| Brain region | Side | Cluster size | MNI coordinates (peak voxel) | | |
|---|---|---|---|---|---|
| | | | x | y | z |
| Delay 0-2 s | | | | | |
| Superior parietal lobe | Left | 68 | -27 | -43 | 62 |
| Supplementary motor area | Bilateral | 86 | 3 | 17 | 46 |
| Inferior frontal gyrus | Right | 81 | 39 | 5 | 18 |
| Middle frontal gyrus | Right | 60 | 42 | 38 | 26 |
| Delay 2-4 s | | | | | |
| Midcingulate cortex | Left | 81 | -9 | 5 | 42 |
| Superior parietal lobe | Left | 334 | -24 | -31 | 62 |
| Middle temporal gyrus | Left | 63 | -54 | -19 | -6 |
| Primary motor area | Bilateral | 74 | 0 | -13 | 66 |
| Inferior frontal gyrus | Right | 57 | 39 | 20 | 42 |
| Middle orbital gyrus | Right | 64 | 39 | 50 | -2 |
| Delay 4-6 s | | | | | |
| Precental/Postcentral gyrus | Left | 462 | -33 | -10 | 58 |
| Visual cortex | Bilateral | 1158 | 0 | -88 | 18 |
| Supplementary motor area | Bilateral | 62 | 0 | -7 | 54 |
| Precentral gyrus | Right | 102 | 30 | -22 | 66 |
| Inferior frontal gyrus | Right | 78 | 51 | 8 | 34 |

**Supplementary Table S2. Summary of searchlight decoding self-/other-attribution during the Delay period**

A threshold at $p < 0.01$ (FWE-corrected at cluster-level with a cluster forming threshold $p < 0.0005$) was set for statistical testing. We applied MVPA to fMRI data in each 2 s time bin (a single scan with TR = 2 s), which was shifted by 6 s later from the actual time considering the HRD (see also Supplementary Fig 3). All clusters larger than 50 voxels are reported.

| | | | | MNI coordinates (peak voxel) | | |
|---|---|---|---|---|---|---|
| Brain region | Side | Cluster size | *T*-value at peak | x | y | z |
| Cycle 3 | | | | | | |
| Self-attribution vs. Other-attribution | | | | | | |
| Cerebellum (VIII A) | Bilateral | 58 | 5.58 | 3 | -73 | -34 |
| Other-attribution vs. Self-attribution | | | | | | |
| Posterior medial frontal cortex | Left | 36 | 3.80 | -12 | 11 | 66 |
| Inferior frontal gyrus (BA44) | Right | 39 | 6.65 | 42 | 5 | 34 |
| Inferior frontal gyrus (BA45) | Right | 36 | 7.80 | 45 | 23 | 42 |
| Inferior parietal lobe | Right | 43 | 3.71 | 54 | -52 | 14 |
| Cycle 4 | | | | | | |
| Self-attribution vs. Other-attribution | | | | | | |
| Cerebellum (VIII A) | Bilateral | 76 | 8.80 | -3 | -61 | -26 |
| Other-attribution vs. Self-attribution | | | | | | |
| Inferior parietal lobe | Right | 68 | 5.95 | 54 | -52 | 14 |

**Supplementary Table S3. Summary of mass-univariate analysis results**

A threshold at $p < 0.001$, corrected for multiple comparisons at the cluster level ($p < 0.05$), was set for statistical testing. We applied mass-univariate analysis to fMRI data corresponding to each of five cycles (shifted by 6 s considering the HRD, see also Supplementary Fig. 4). We could not find a significant cluster for the cycle 1, 2 or 5 at the same threshold.

| Participant | Slope | SD |
|---|---|---|
| 1 | 6.33 | 1.58 |
| 2 | 5.41 | 1.91 |
| 3 | 4.05 | 2.05 |
| 4 | 4.99 | 1.79 |
| 5 | 4.23 | 2.62 |
| 6 | 8.37 | 1.11 |
| 7 | 6.26 | 1.45 |
| 8 | 6.37 | 1.16 |
| 9 | 5.6 | 1.35 |
| 10 | 4.65 | 1.46 |
| 11 | 5.65 | 1.59 |
| 12 | 4.41 | 2.92‡ |
| 13 | 3.49† | 2.13 |
| 14 | 3.15† | 1.70 |
| 15 | 1.74† | 2.55 |
| 16 | 5.11 | 2.90‡ |
| 17 | 3.27† | 3.02‡ |
| 18 | 4.29 | 3.68‡ |

**Supplementary Table S4. Slope value of regression line and standard deviation (SD) of rating scores in self 0% condition.**

The participants under the horizontal dotted line (i.e., Participants 12 ~ 18) were eliminated from the analysis due to low slope values (< 3.89, marked with †) and/or large SDs (> 2.89, marked with ‡).
